# SERPINA3 and NDRG1 are critical diagnostic immune genes associated with macrophages in preeclampsia

**DOI:** 10.64898/2026.02.09.704892

**Authors:** Zhuna Wu, Shihong Chen, Weihong Chen, Yajing Xie, Zhimei Zhou, Li Huang, Liying Sheng, Yueli Wang, Binbin Chen, Congmei Yang, Yumin Ke

## Abstract

**Objective:** The immune system plays a role in the occurrence and progression of numerous pregnancy complications, particularly preeclampsia (PE). This study aims to identify critical immune biomarkers via machine learning and assess their predictive ability.

**Methods:** Gene expression data were retrieved from the GEO database, while immune-related genes were obtained from the ImmPort repository. Differential expression analysis was then conducted to identify immune genes associated with PE. Different immune-related genes (DIRGs) were subjected to functional and pathway enrichment analysis. We adopted protein-protein interaction (PPI) networks for exploring the connections among various DIRGs and integrated two machine-learning to pinpoint candidate biomarkers in PE. Diagnostic performance was assessed via ROC curve analysis, with predictive accuracy further quantified using nomogram calibration. Findings were validated through integrated computational and experimental analyses. In silico validation utilized additional GEO datasets, while experimental confirmation involved qRT-PCR and IHC assessment of placental tissues. We developed a nomogram to predict PE utilizing two immune-related genes. Cellular composition was inferred from transcriptomic data using CIBERSORT deconvolution..

**Results:** We identified 66 differentially expressed genes (DEGs) and 10 DIRGs between PE pregnancies and normotensive pregnancies. The GO analyses revealed that the DIRGs were enriched in gonadotropin secretion, the regulation of gonadotropin secretion, and the regulation of endocrine processes. Functional annotation revealed enrichment in cytokine and neuroactive ligand-receptor pathways. SERPINA3 and NDRG1 emerged as top-performing biomarkers (training AUCs: 0.812 and 0.866; external validation: 0.795 and 0.781), with overexpression validated in clinical specimens. Both genes inversely regulated M2 macrophage abundance (P < 0.05).

**Conclusion:** PE is fundamentally an immune-mediated disorder. SERPINA3 and NDRG1 can be identified as key immune genes associated with M2 macrophages, and these findings provide novel perspectives for the diagnosis and pathogenesis of PE.

## Introduction

Preeclampsia (PE) is a serious disease during pregnancy and is a prominent cause of neonatal and maternal mortality and morbidity^[1]^. PE is a multifaceted disorder characterized by the abrupt development of high blood pressure (after 20 weeks of pregnancy) along with at least one additional related complication, such as proteinuria, maternal organ impairment, or dysfunction of the uteroplacental unit. Globally, approximately four million women are diagnosed with PE annually, resulting in the fatality of more than 70,000 women and 500,000 infants^[2]^. On a global scale, more than 300 million women and children are projected to face increased susceptibility to long-term health issues as a result of past exposure to PE^[3]^. Given the current lack of screening before the occurrence of PE, it is essential to pinpoint innovative diagnostic biomarkers to identify potential PE earlier and therapeutic targets to improve outcomes for fetuses and mothers with PE.

Gestational immune imbalance cascades from maternal systemic inflammation to impaired fetal immune maturation and persistent postnatal immunological alterations. Multiple innate and adaptive immune cells and factors have been linked to the pathogenesis of PE, where oxidative stress is connected to the activation of the maternal inflammatory response^[4]^. The innate immune system, which involves components such as complement proteins^[5]^, neutrophils^[6]^, macrophages^[7]^, and NK cells^[8]^, serves to defend both the mother and fetus against infections while also playing a role in the formation of the maternal‒fetal interface. Adaptive immune responses mediated by T and B cells can target pathogens as well as auto and alloantigens. These responses are distinguished by the development of immune memory, which boosts the immune reaction upon future exposure to the same antigens. Notably, PE occurs more frequently in initial pregnancies than in subsequent pregnancies^[9]^.

Hence, the objective of this study was to identify novel DIRGs associated with PE samples for the purpose of identifying diagnostic immune biomarkers through the utilization of bioinformatics approaches. Further efforts were undertaken to validate the DIRGs among placental samples from the normotensive pregnancy and PE groups. Additionally, we investigated the potential correlation between novel DIRGs and immune cells to foster additional research on the pathogenesis of PE in this field.

## Methods and Materials

### Data acquisition and preprocessing

Publicly available transcriptomic datasets related to preeclampsia (PE) were retrieved from the Gene Expression Omnibus (GEO) database. The dataset GSE75010 (accessed on 13/03/2024) was designated as the training cohort, while GSE54618, GSE74341, and GSE147776 (accessed on 13/03/2024) were merged to form an independent validation cohort. Detailed cohort specifications, including platform annotations and sample sizes, are documented in Table 1. Preprocessing comprised background subtraction followed by between-array normalization using limma (R v4.1.3). Probes were mapped to gene symbols based on platform annotation files, and for genes represented by multiple probes, the average expression value was retained. Batch effects among the validation datasets were mitigated using the “sva” package.

**Table 1:**
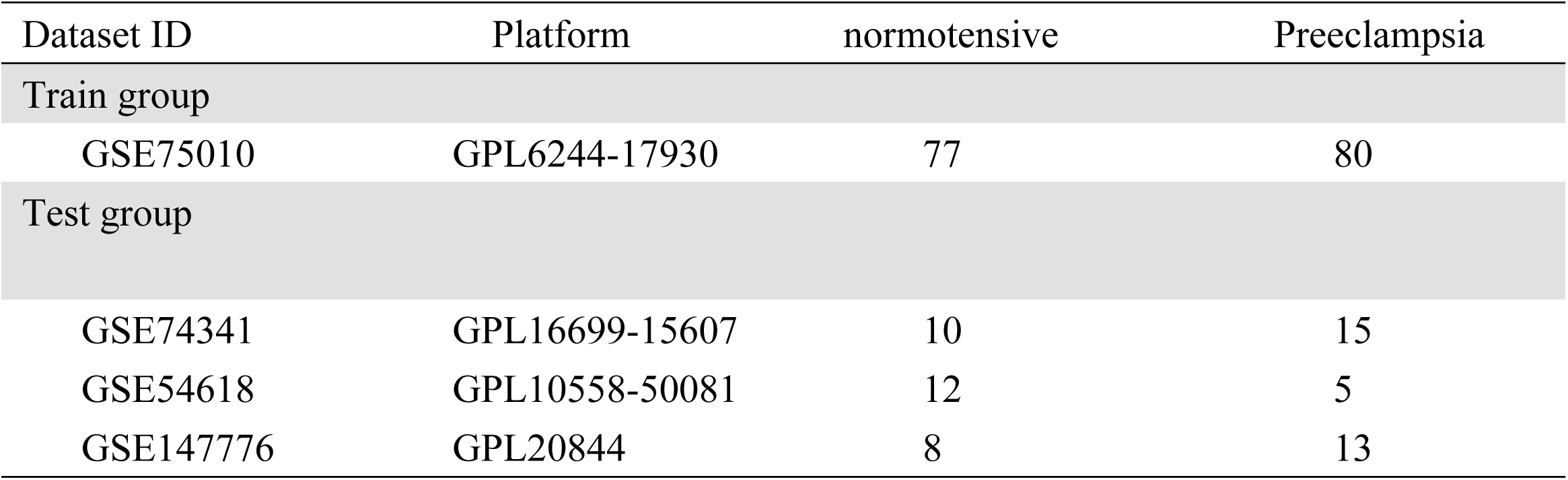
mRNA expression profiles related to PE from the GEO database.

| Dataset ID | Platform | normotensive | Preeclampsia |
| --- | --- | --- | --- |
| Train group |  |  |  |
| GSE75010 | GPL6244-17930 | 77 | 80 |
| Test group |  |  |  |
| GSE74341 | GPL16699-15607 | 10 | 15 |
| GSE54618 | GPL10558-50081 | 12 | 5 |
| GSE147776 | GPL20844 | 8 | 13 |

### Identification of differentially expressed immune-related genes

To identify differentially expressed genes (DEGs) associated with PE, we performed comparative transcriptomic analysis between PE and normotensive samples in the GSE75010 dataset using the ‘limma’ R package. Statistical significance thresholds comprised a fold-change cutoff of 1.5 (|log₂FC| > 0.585) and multiple-testing corrected P values < 0.05. The immune gene reference set was curated from ImmPort (https://www.immport.org, accessed March 2024)(Supplementary Table S1). The intersection between DEGs and IRGs yielded differentially expressed immune-related genes (DIRGs) for further analysis.

### Ontology and Pathway Profiling

Functional interpretation employed over-representation analysis to delineate statistically enriched biological themes and molecular pathways. This included Gene Ontology (GO) analysis covering three categories and Kyoto Encyclopedia of Genes and Genomes (KEGG) analysis, implemented through the clusterProfiler suite of R packages (“clusterProfiler”, “enrichplot”, “org.Hs.eg.db”, and “DOSE”). We utilized the “ggplot2” package in R to visualize the enrichment results. A significance level of p < 0.05 was used to indicate significant enrichment.

### Protein–protein interaction (PPI) network construction and Analysis

Protein-protein interaction (PPI) networks were reconstructed using the STRING database (version 11.5, https://string-db.org). The 10 DIRGs were batch-queried through the multiple protein analysis module with organism restricted to Homo sapiens. Interactions lacking experimental validation or database curation were excluded, and disconnected network nodes were pruned to retain the largest connected component. The filtered interaction matrix was exported and imported into Cytoscape (version 3.10.0) for topological analysis. Hub gene prioritization was performed using the cytoHubba plugin, applying the Maximal Clique Centrality (MCC) algorithm—a local network topology metric that identifies nodes participating in the maximum number of k-cliques—to rank genes by their topological significance within the immune-related subnetwork.

### Machine learning–based biomarker selection

Candidate biomarker selection was initiated by filtering DIRGs based on effect size (|log₂ fold change| ≥ 0.585) and correlation strength (Spearman’s ρ > 0.5). Feature dimensionality reduction subsequently employed two complementary machine learning paradigms: (i) LASSO regression (glmnet package), utilizing L1-norm regularization (λ= 0.01–1.0 via 10-fold cross-validation) to shrink less informative feature coefficients toward zero, thereby preventing model overfitting and ensuring predictor parsimony; and (ii) mSVM-RFE (e1071 package), employing a linear kernel SVM classifier with recursive backward elimination to systematically discard features exhibiting minimal weight contributions to the hyperplane decision boundary^[11]^. Consensus genes retained by both algorithms were designated as high-confidence diagnostic biomarkers..

### Diagnostic model development and validation

Discriminatory accuracy was quantified by area under the ROC curve (AUC) using the pROC R package. The area under the curve (AUC) was calculated to quantify discriminatory power in both training and validation cohorts. A nomogram integrating selected biomarkers was constructed with the “rms” package to visualize individual and combined predictive contributions. Calibration curves were plotted to assess model accuracy.

### Clinical sample collection

This study was approved by the Research Ethics Committee of the Second Affiliated Hospital of Fujian Medical University (Approval No. 2024-321). All participants provided written informed consent. Between December 2020 and May 2024, placental tissues were collected from 38 PE patients and 40 normotensive controls undergoing cesarean or vaginal delivery at the same institution. Tissues were either paraffin-embedded for immunohistochemistry or snap-frozen for RNA extraction.

### In Situ Protein Detection and Localization

Standard IHC procedures were adapted and optimized for placental tissue analysis^[12]^. In brief, tissue sections were deparaffinized, subjected to antigen retrieval, and incubated overnight at 4°C with primary antibodies against SERPINA3 (Bioss, Beijing) and NDRG1 (Affinity, USA). Following secondary antibody incubation and DAB development, staining was scored independently by two pathologists blinded to clinical data. Immunohistochemical scoring employed a multiplicative index (intensity × distribution) with dichotomous stratification (low: <6; high: ≥6). Chromogenic intensity was graded 0–3 (negative, weak, moderate, strong) and cellular coverage 1–3 (<33%, 33–66%, >66%). Histopathological diagnoses were independently verified by two specialist pathologists.

### Transcript Quantification by qPCR

Snap-frozen placental specimens underwent total RNA purification via the TRIzol method (Beyotime, China). cDNA was synthesized with a PrimeScript RT kit (TaKaRa, Japan). qRT - PCR was performed in triplicate using SYBR Green assays on a QuantStudio 5 system (Applied Biosystems). GAPDH served as the endogenous control, and relative expression was calculated via the 2−ΔΔCT method. Primer sequences are listed below:

#### GAPDH

Forward: 5′-GTCTCCTCTGACTTCAACAGCG-3′,

Reverse: 5′-ACCACCCTGTTGCTGTAGCCAA-3′.

#### SERPINA3

Forward: 5′-TCTGGAGTTCAGAGAGATAGGTGAG-3′,

Reverse: 5′-TGGAGAAGTATGTCGTTCAGGTTAT-3’.

#### NDRG1

Forward: 5′-AGAGGCAATGACTCGTTACCTG-3′,

Reverse: 5′-GCACAATAGTTTCCCATCCGTA-3’.

### Immune cell infiltration analysis in PE

Cellular deconvolution of bulk transcriptomic data was performed using the CIBERSORT algorithm (version 1.06) with the LM22 leukocyte gene signature matrix, which comprises 547 genes distinguishing 22 hematopoietic cell phenotypes^[14]^. Samples with CIBERSORT output P < 0.05 were retained for downstream analysis to ensure reliable fraction estimation. Differential abundance analysis of immune cell subsets between PE and control groups was conducted using the Wilcoxon rank-sum test with Benjamini-Hochberg false discovery rate (FDR) correction (q < 0.05 considered significant). Associations between SERPINA3/NDRG1 expression levels and immune cell proportions were evaluated using Pearson’s product-moment correlation, with resultant correlation matrices visualized via the corrplot and ggplot2 R packages.

### Statistical analysis

Data analysis was carried out in R version 4.1.3. For comparative analyses, we used Mann-Whitney U tests for group comparisons and unpaired t-tests for continuous variables. Feature selection involved LASSO regression and SVM-RFE algorithms, while diagnostic evaluation utilized ROC curve analysis. Correlation analyses employed Pearson’s method.

## Results

### Study procedure

As depicted in Figure 1, our analytical approach began with RNA-seq data acquisition from GEO. We then identified differentially expressed genes and intersected them with immune gene annotations to obtain differentially expressed immune-related genes for PE. Comprehensive bioinformatic interrogation included: (i) functional enrichment against Gene Ontology and KEGG databases; (ii) PPI network reconstruction with hub gene identification; and (iii) diagnostic biomarker selection through penalized regression (LASSO) and recursive feature elimination (SVM-RFE). Classifier performance was evaluated by receiver operating characteristic curves and prospectively validated in a composite validation cohort (GSE54618+GSE74341+GSE147776). The compositional patterns of LM22 in PE were calculated via the CIBERSORT algorithm. Additionally, correlation analysis was conducted between diagnostic immune-related biomarkers and infiltrating immune cells.

**Figure 1:**
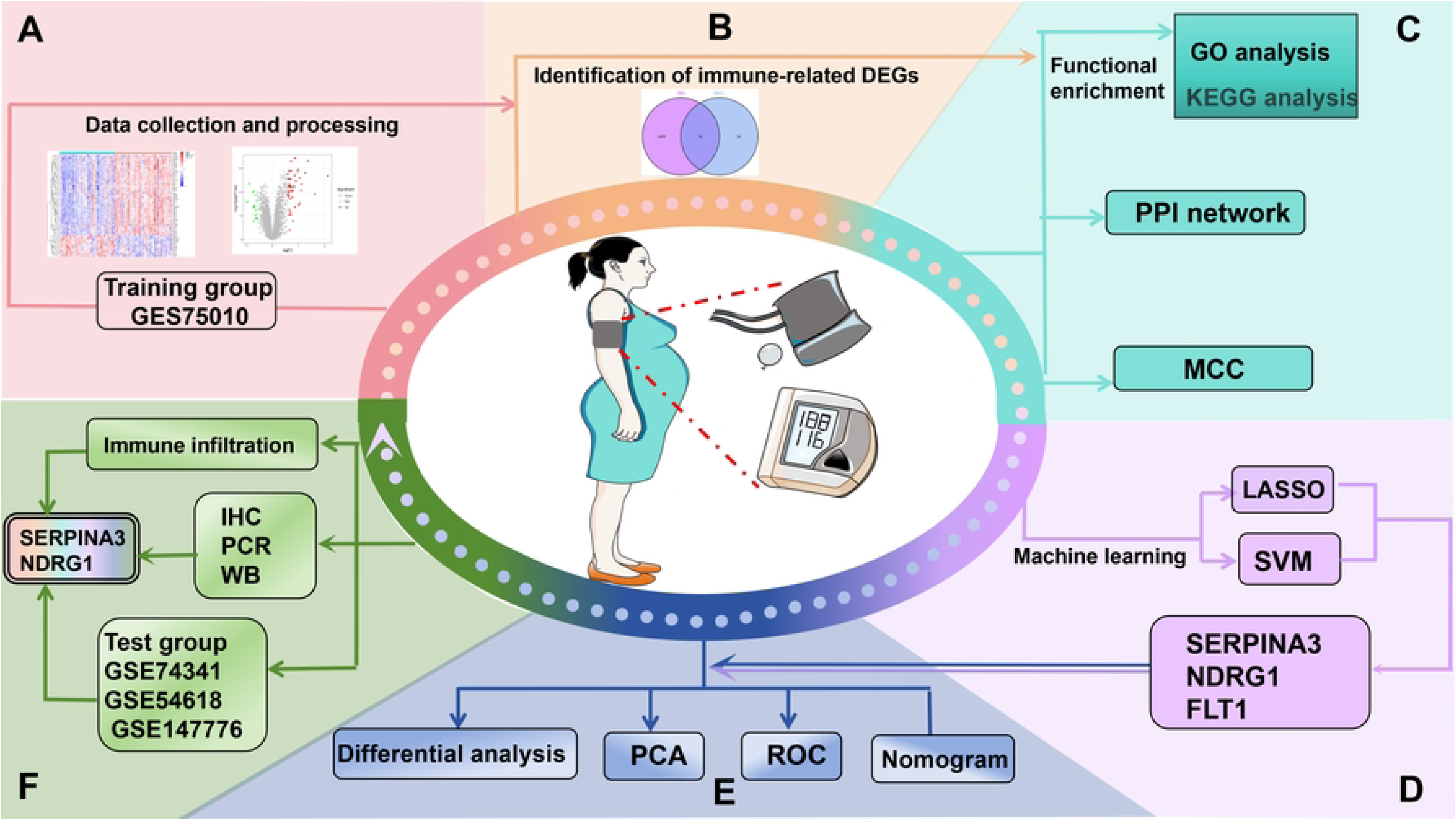
Workflow diagram of this research.

### Discovery of PE-Associated Immune Genes

Comparative transcriptomics using empirical Bayes moderation (limma) revealed 66 significant DEGs in the discovery cohort (GSE75010), with effect size threshold of 1.5-fold change and multiple-testing correction (adj. P < 0.05). The transcriptional landscape was dominated by upregulation (52/66, 78.8%). Filtering against curated immune annotations identified a core set of 10 DIRGs exhibiting coordinated immune dysregulation (Figure 2C). Among the 10 DIRGs, OPRK1 expression was lower in the PE group, whereas 9 DIRGs were significantly upregulated in the PE group.

**Figure 2:**
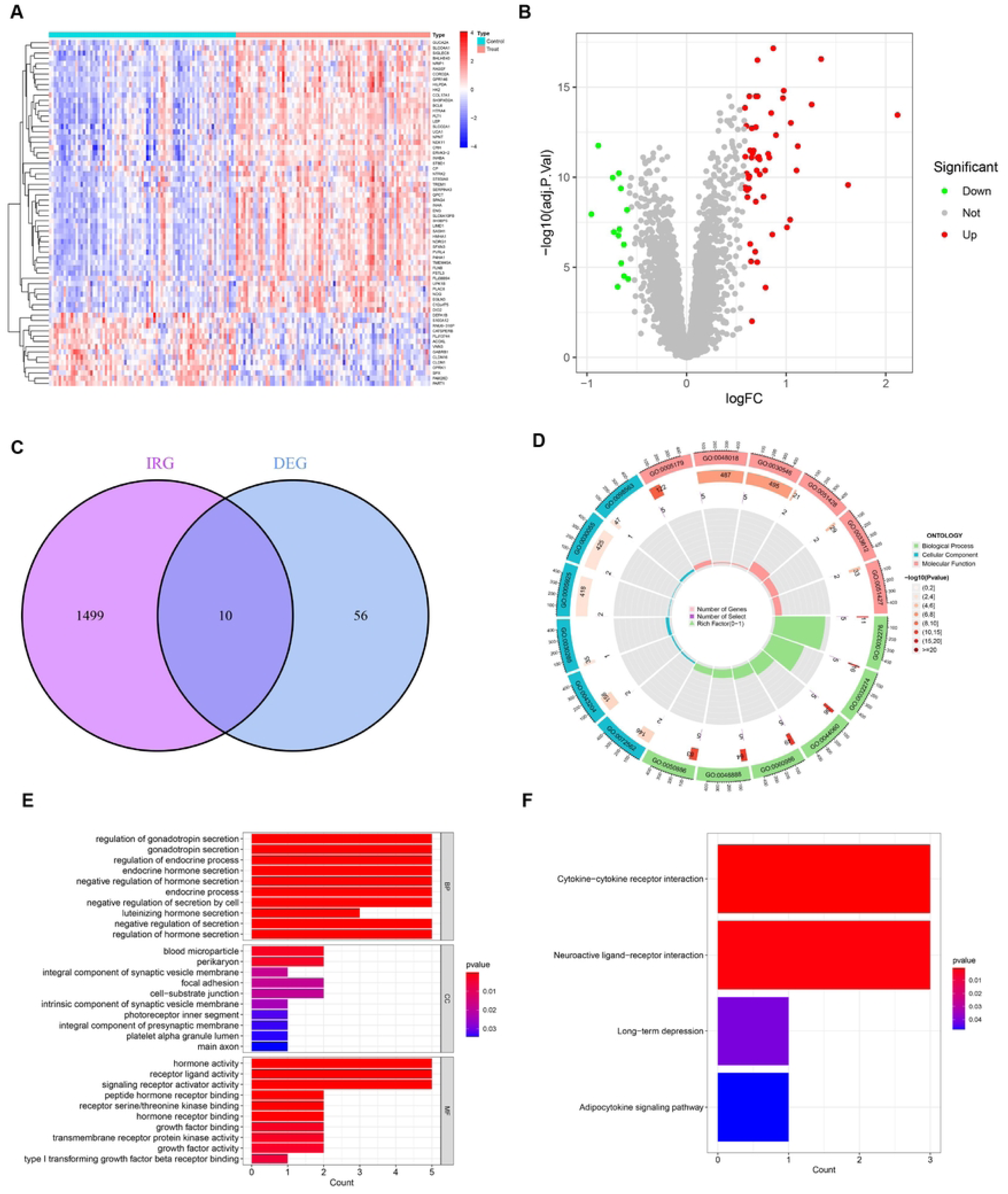
Identification and function of DIRGs. (A)Heatmaps of the expression levels of 66 DEGs between PE tissue and normotensive control pregnancy samples from the GEO database are visualized. The genes are identified by their names in the row annotations, while the column annotations are the sample IDs, which are not displayed in the plots. The color gradient, ranging from red to blue, signifies the expression levels from high to low in the heatmaps. (B)The volcano plots illustrate 66 DEGs between PE tissue and normotensive control pregnancy samples. In these plots, red dots indicate genes that are upregulated, green dots denote genes that are downregulated, and black dots represent genes that do not show differential expression. (C)The intersection of DEGs in the GSE75010 dataset and IRGs downloaded from ImmPort contains 10 DIRGs. (D)Circle plot of GO analysis for 10 DIRGs. (E) bar graph of GO analysis for 10 DIRGs. (F)KEGG annotation of DIRGs.

### Functional enrichment (GO and KEGG)

Systems-level pathway annotation was performed to delineate the biological processes and signaling cascades enriched within the DIRG signature. These genetic biological processes (BP) are predominantly focused on the regulation of gonadotropin secretion and the regulation of the endocrine process. The cellular components (CCs) of the DIRGs are located primarily in blood microparticles, perikaryons, and focal adhesions. The molecular functions (MFs) of the DIRGs were associated primarily with receptor ligand activity, signaling receptor activator activity, and peptide hormone receptor binding (P < 0.05, Figure 2D-E, Supplemental Figure S1). In addition, KEGG enrichment analysis suggested that the 10 DIRGs were predominantly involved in cytokine‒cytokine receptor interactions, neuroactive ligand‒receptor interactions, and adipocytokine signaling pathways (P <0.05, Figure 2F, Supplemental Figure S2). These results indicate that PE is strongly associated with immunity.

### Development of the PPI and Hub Genes network

The Search Tool for the Retrieval of Interacting Genes/Proteins (STRING) database (https://string-db.org/) was used to examine the interactions of 10 immune genes, which produced protein–protein interaction (PPI) networks with hidden disconnected nodes in the network (see Figure 3A). The cytoHubba plugin within the Cytoscape software was subsequently utilized to cluster the genes in the network. The eight nodes associated with the MCC were identified and grouped (Figure 3B). All the genes presented increased expression (Figure 3C, 3D).

**Figure 3:**
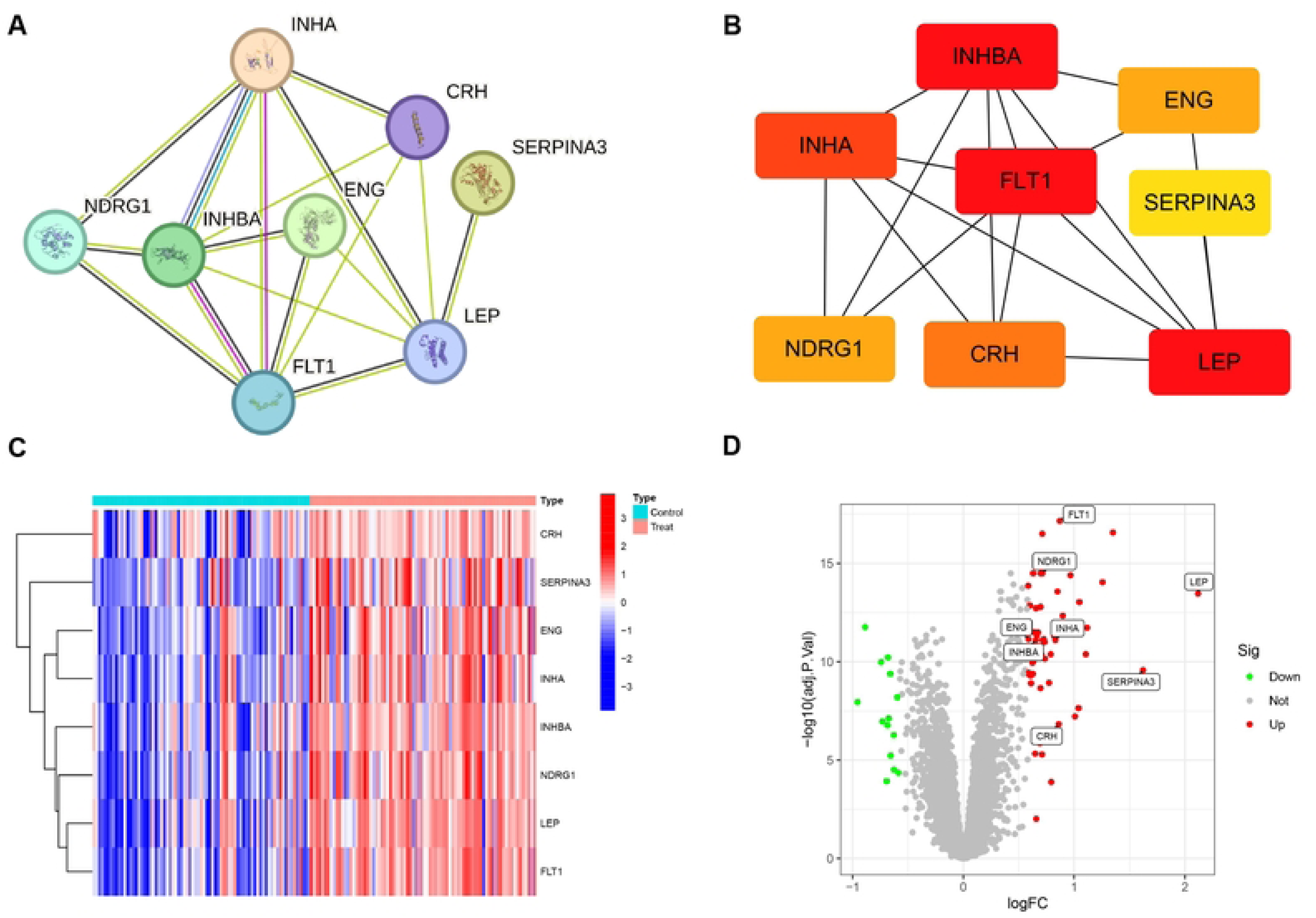
Correlation between DIRGs and hub genes. (A)Using the STRING tool to construct PPI networks, exploring eight DIRGs binding protein interactions. (B)Eight hub genes obtained by the MCC algorithm. (C)Heatmaps of the expression levels of eight hub genes between PE tissue and normotensive control pregnancy samples. (D)The volcano plots illustrate eight hub genes between PE tissue and normotensive control pregnancy samples.

### Associations among the expression levels of DIRGs in PE

Inter-gene correlation networks were constructed to elucidate transcriptional co-regulation patterns among the 8 DIRGs. Pairwise correlation coefficients were computed using Spearman’s rank correlation (|r| > 0.4), with subsequent visualization implemented through the tidyverse and corrr R packages to generate both network topology maps (Figure 4A) and correlation matrices (Figure 4B). Stringent filtering criteria (correlation coefficient |r| > 0.6, adjusted P < 0.001) identified five robustly correlated gene pairs, which were further illustrated through scatter plot matrices (Figure 4C). Notably, NDRG1 exhibited strong positive correlations with FLT1 (r = 0.78), INHA (r = 0.72), LEP (r = 0.69), and INHBA (r = 0.65), suggesting potential co-regulatory mechanisms or functional interdependencies among these genes in PE pathophysiology.

**Figure 4:**
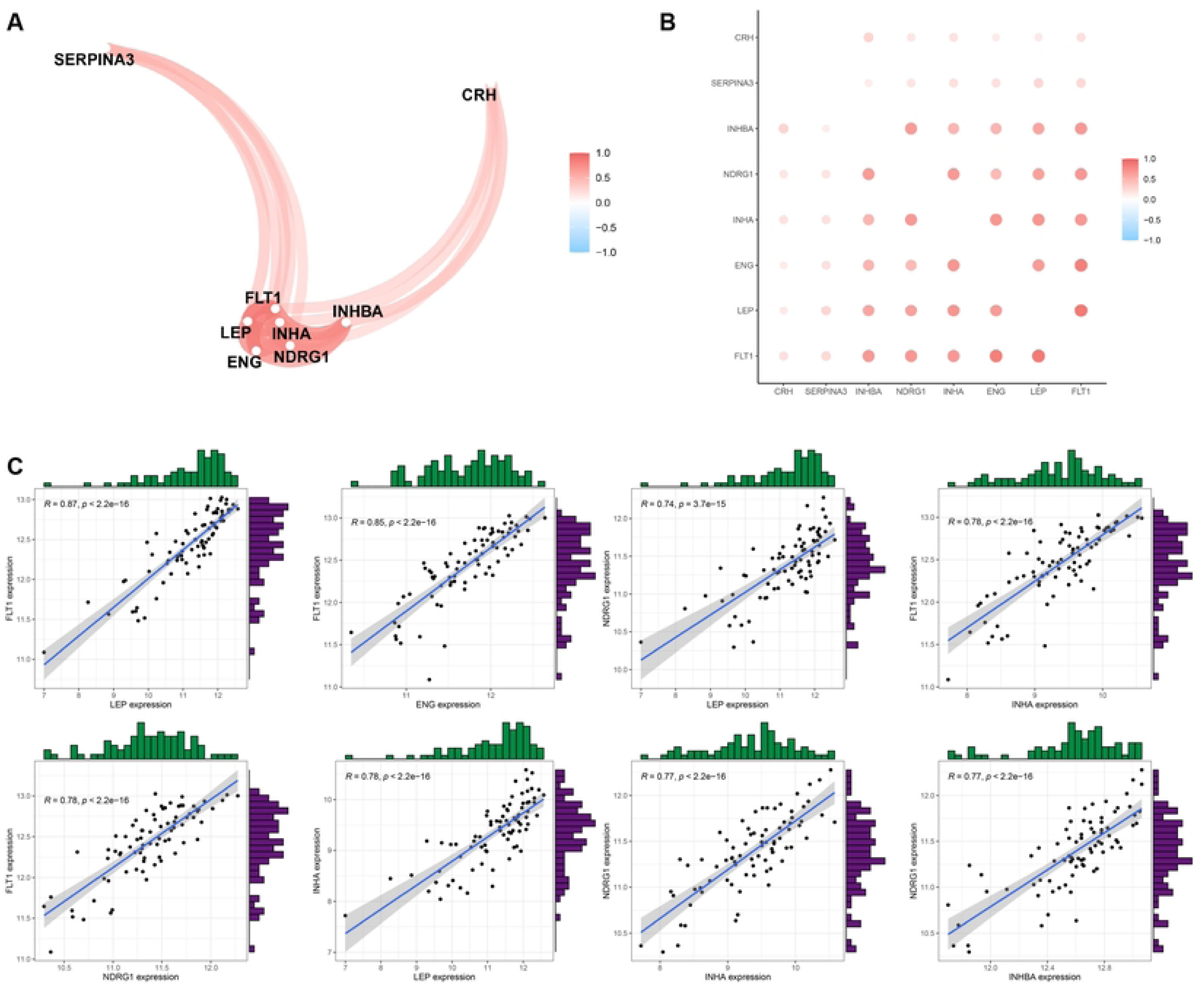
Correlation analysis on DIRGs. (A)Co-expression network map of DIRGs. (B) Correlation plot of DIRGs. (C)Scatter plot of some highly correlated DIRGs.

### Development of a prediction model for PE

To identify robust diagnostic biomarkers for PE, we implemented a dual-machine learning strategy integrating Least Absolute Shrinkage and Selection Operator (LASSO) regression with multiple Support Vector Machine-Recursive Feature Elimination (mSVM-RFE). LASSO regression, which applies L1 regularization to minimize overfitting and select optimal features, identified a panel of 3 DIRGs (SERPINA3, NDRG1, and FLT1) from the initial 8 differentially expressed immune-related genes (Figure 5A, 5B). Concurrently, mSVM-RFE analysis, which iteratively eliminates less informative features based on SVM-derived weights, narrowed the candidate pool to 7 DIRGs comprising FLT1, SERPINA3, INHBA, NDRG1, INHA, CRH, and ENG (Figure 5C, 5D). The intersection of these two algorithmic outputs yielded a consensus set of 3 candidate biomarkers (SERPINA3, NDRG1, and FLT1) with high diagnostic potential (Figure 5E)..

**Figure 5:**
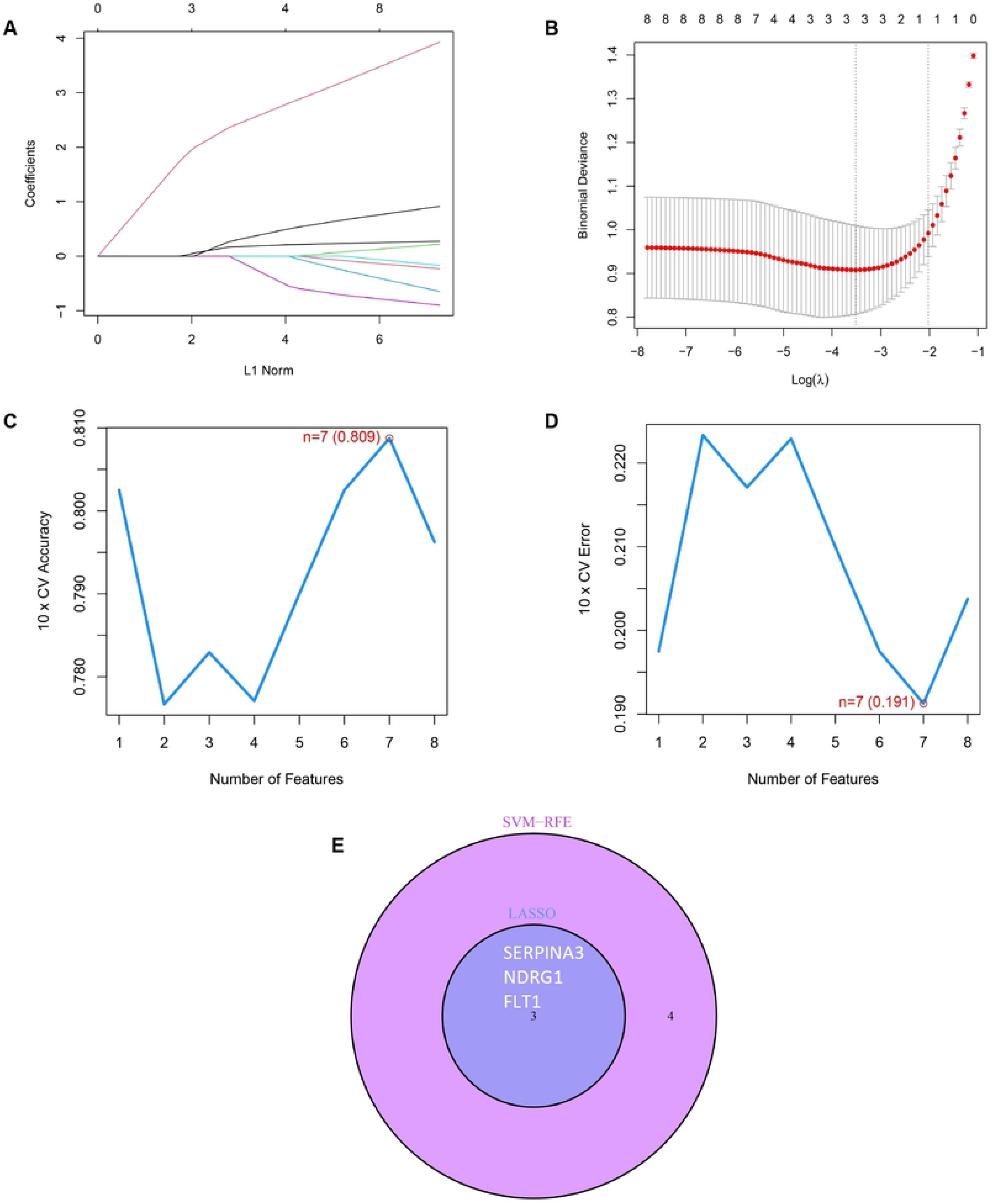
Development of a prediction model for PE. (A)The LASSO regression coefficient profiles of the 8 DIRGs depict the changing trajectory of each DIRG through a curve. (B)The LASSO Cox regression model was used to plot partial likelihood deviance versus log (l). (C)The curve of the total within sum of squared error curve under the corresponding cluster number k reached the “ elbow point ” when k = 7. (D)The curve representing average silhouette width for the corresponding cluster number k attained its peak at k = 7. (E)The Venn diagram illustrates the three diagnostic markers shared by the LASSO and SVM-RFE algorithms.

### Additional examination of the three key DIRGs

The consensus biomarkers (SERPINA3, NDRG1, and FLT1) emerging from the convergence of LASSO and mSVM-RFE algorithms underwent comprehensive downstream validation. Genomic mapping revealed distinct chromosomal localizations for these genes: SERPINA3 at 14q32.13, NDRG1 at 8q24.22, and FLT1 at 13q12.2 (Figure 6A). Principal component analysis (PCA) demonstrated that the combined expression profiles of these three genes achieved clear segregation between PE and normotensive control samples, underscoring their discriminatory power in distinguishing pathological from physiological pregnancies (Figure 6B). Within the training cohort (GSE75010), all three biomarkers exhibited significantly elevated transcript levels in PE-derived placental tissues relative to gestational age-matched controls (Figure 6C). Receiver operating characteristic (ROC) curve analysis quantified their individual diagnostic performances, with NDRG1 demonstrating the highest discriminatory accuracy (AUC = 0.866), followed by FLT1 (AUC = 0.886) and SERPINA3 (AUC = 0.812), collectively surpassing the predictive threshold for clinical utility (Figure 6D). A risk-related nomogram for PE was subsequently constructed (Figures 6E and 6F), which can serve as a predictor of the ability of the risk score to distinguish between PE and normotensive control pregnancies.

**Figure 6:**
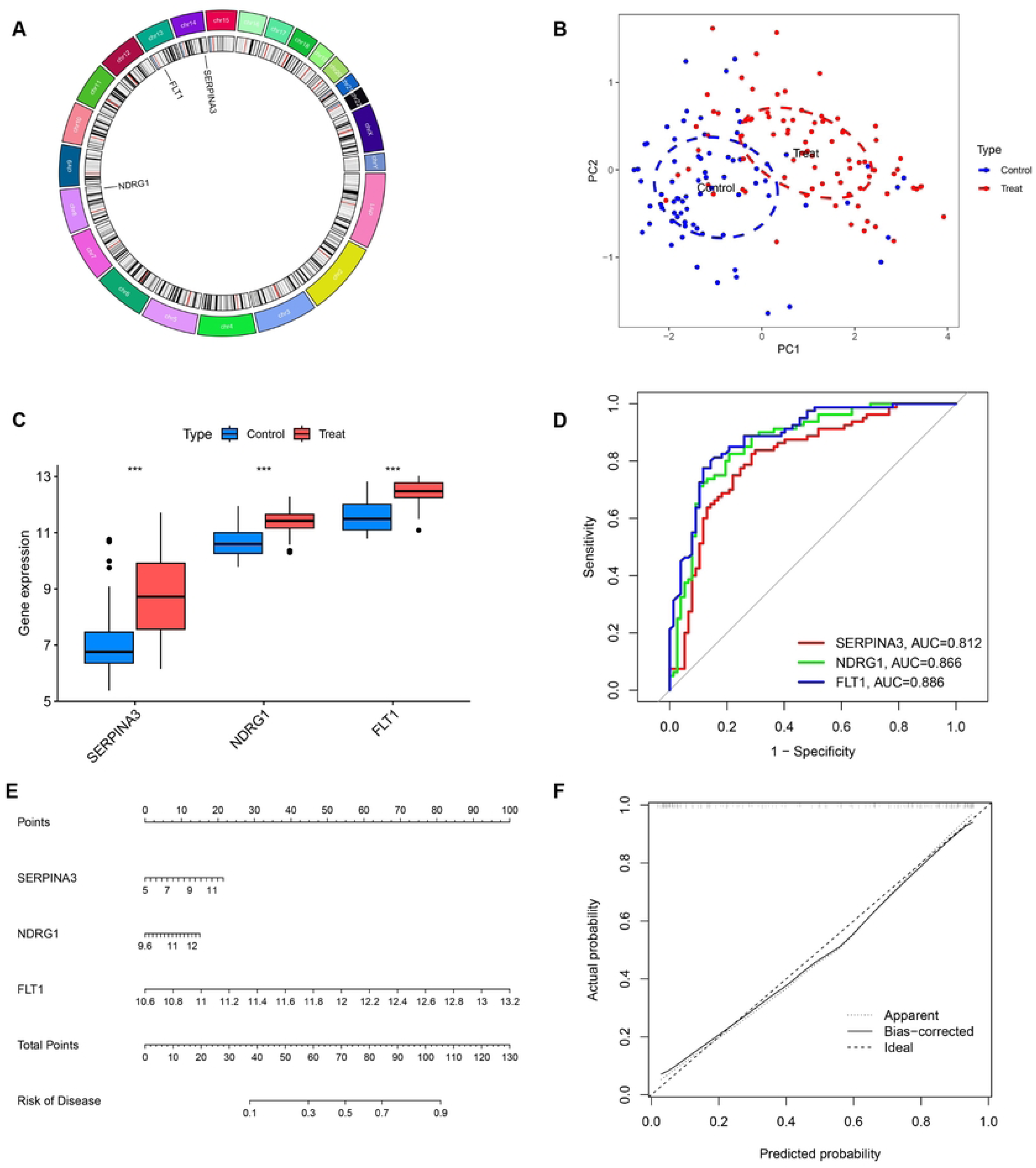
Further analysis of three key DIRGs. (A)The chromosomal locations of three key DIRGs. (B)Principal component analysis clearly distinguishes PE and normotensive control pregnancy using the three aforementioned genes. (C)The GSE75010 datasets reveal the relative expression levels of three key DIRGs between PE and normotensive control pregnancy. (D)ROC curves validated the performances of three key DIRGs for the prediction of PE in GSE75010 datasets. (E) Diagnostic Nomo plot of three key DIRGs. (F) Calibration curve of a model composed of three key genes.

### Validation of the three key DIRGs

The predictive efficacy of these three DIRGs was evaluated in the test cohort (GSE54618, GSE74341, and GSE14776). As depicted in Figure 7A, the expression of SERPINA3, NDRG1, and FLT1 was upregulated in PE. Furthermore, strong discerning capacity was validated in the test cohort, with an AUC of 0.798 for SERPINA3 and an AUC of 0.781 for NDRG1 (Figure 7B), indicating that the SERPINA3 and NDRG1 genes have greater diagnostic capacity.

**Figure 7.**
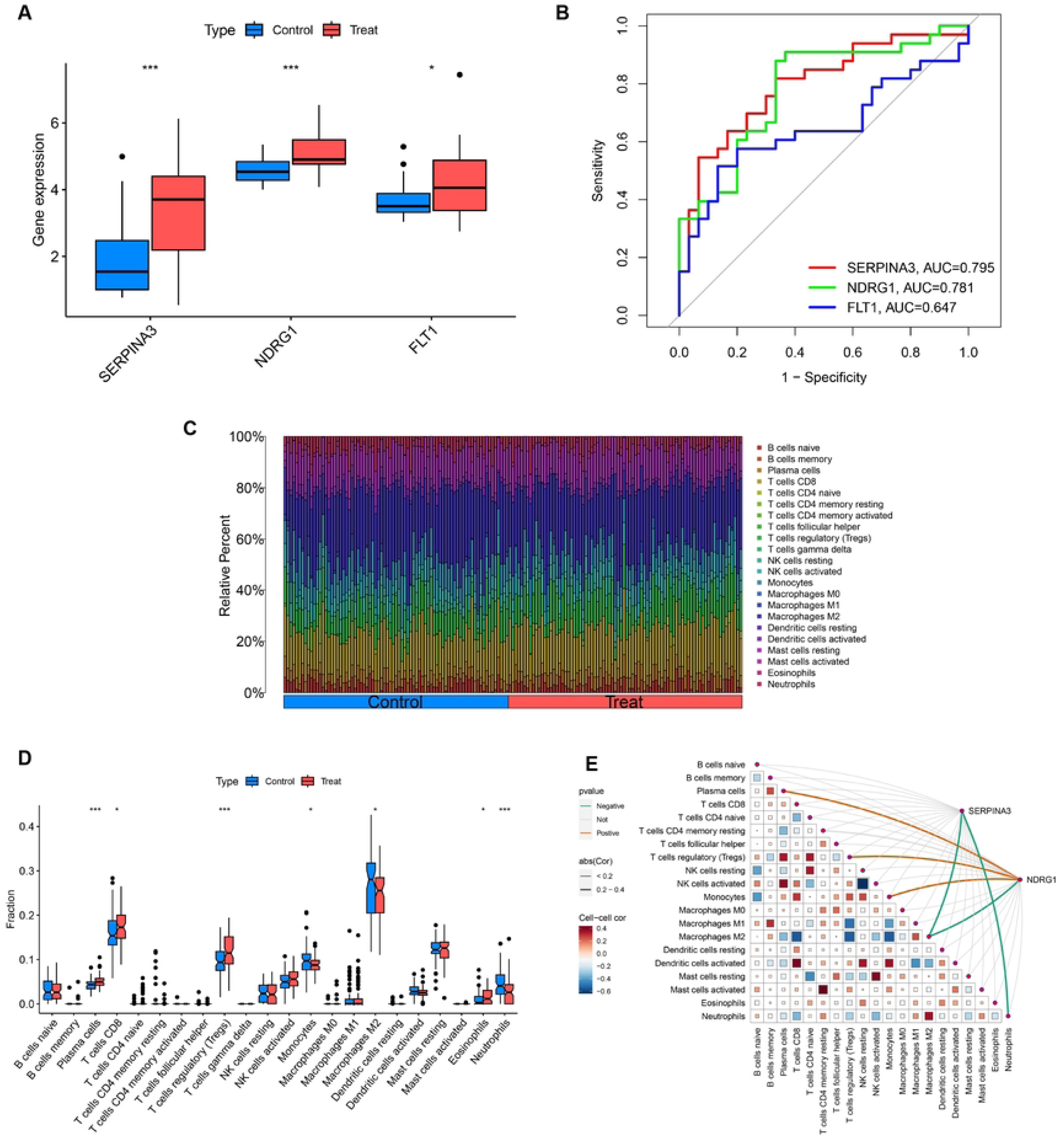
Validation of the three key DIRGs and immune cell infiltration distribution. (A)The relative expression level of three key DIRGs between PE and normotensive control pregnancy from GSE54618, GSE74341, and GSE14776 datasets. (B)ROC curves validated the performances of three key DIRGs for the prediction of PE in GSE54618, GSE74341, and GSE14776 datasets. (C)Bar plot illustrating the distribution of 22 immune cell subtype proportions between PE tissue and normotensive control pregnancy samples. (D)Differences in the infiltrating immune cells between the PE and normotensive control pregnancy group. (E) Correlation between SERPINA3, NDRG1, and infiltrating immune cells in PE.

### SERPINA3 and NDRG1 are associated with the distribution of immune cells

To gain deeper insight into the relationship between immune cell infiltration and PE, we used the CIBERSORT algorithm to determine the relative proportions of 22 types of immune cells in the control and PE samples (Figure 7C). We then compared immune cell infiltration between PE and normotensive control samples. Plasma cells, CD8+ T cells, regulatory T cells (Tregs), and eosinophils were significantly more abundant in PE patients, whereas the infiltration levels of monocytes, M2 macrophages, and neutrophils were significantly lower in PE patients (Figure 7D). The immunomodulatory roles of SERPINA3 and NDRG1 were investigated through correlation analysis with tissue-resident immune populations. SERPINA3 exhibited a negative association with neutrophils and M2 macrophages. NDRG1 was positively associated with plasma cells, Tregs, and monocytes. Conversely, NDRG1 was negatively correlated with M2 macrophages (Figure 7E). Collectively, these correlations between SERPINA3/NDRG1 expression and specific immune cell subsets reinforce the potential role of these genes in modulating the immune microenvironment of PE.

### Clinically validate diagnostic immune biomarkers

We assessed the expression of SERPINA3 and NDRG1 across PE and Non-PE tissues via IHC and revealed that high expression of SERPINA3 and NDRG1 was associated with PE (Figure 8A, 8B; p < 0.05). For further validation, Experimental verification via qRT-PCR aligned with bioinformatic predictions, demonstrating significant overexpression of SERPINA3 and NDRG1 in the disease cohort (Figure 8C; P < 0.05). The results mentioned above suggest that the SERPINA3 and NDRG1 genes have greater diagnostic capacity.

**Figure 8:**
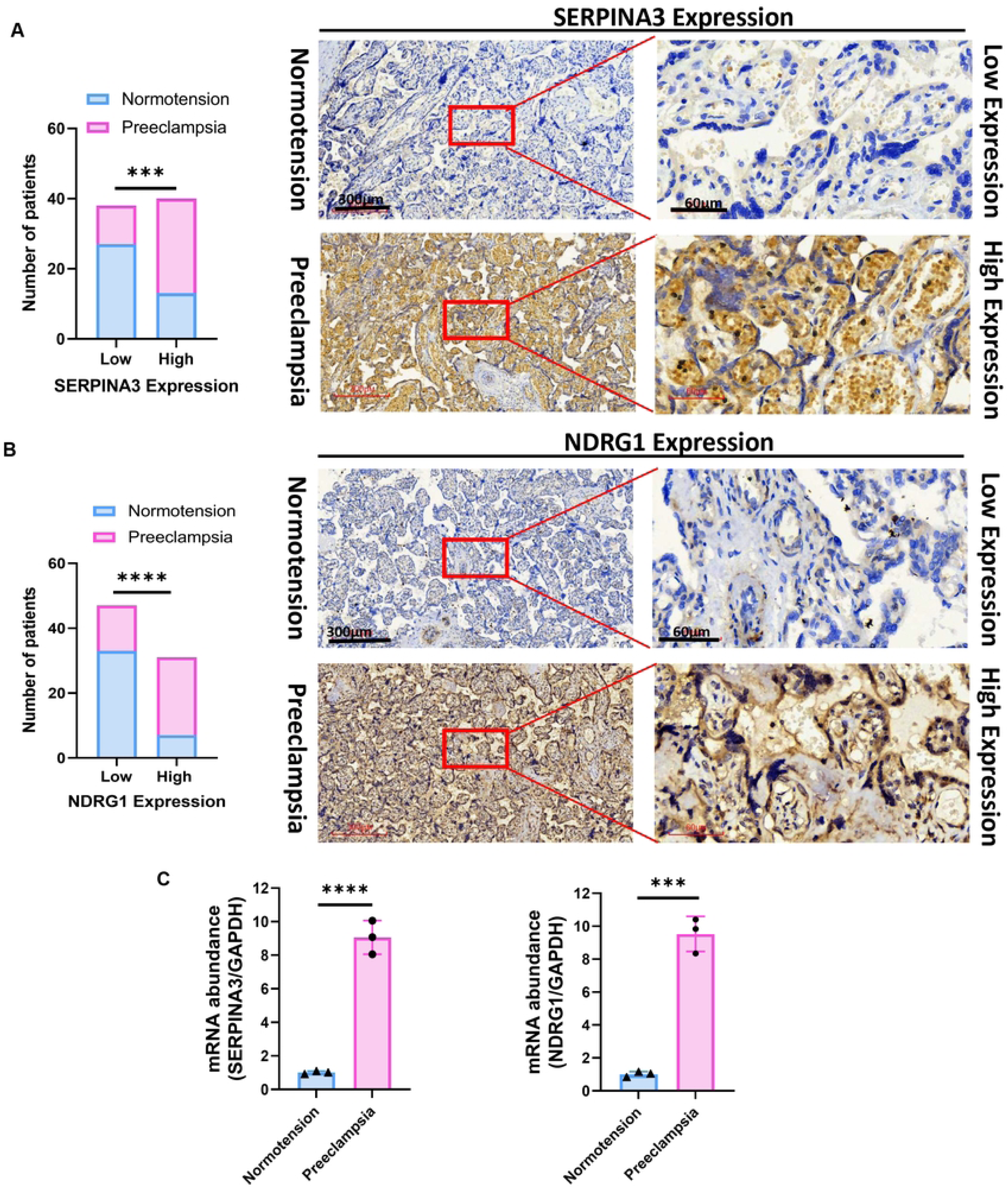
Clinically validate diagnostic immune biomarkers. (A, B) Significantly high SERPINA3 and NDRG1 expression was observed in PE tissues when compared with normotension specimens (normotension = 40; PE = 38). Representative images (×40 and ×200) of IHC staining for SERPINA3 and NDRG1 in 38 PE and 40 normotensive patients (high expression vs. low expression). (C) qRT-PCR of SERPINA3 and NDRG1 expression in PE placental tissue when compared with normotension specimens. Scale bars are shown. ***p < 0.001, ****p < 0.0001. p-values were calculated by chi-square tests.

## Discussion

PE is diagnosed only when the placenta is or has recently been present, and it is categorized into preterm (delivery < 37 weeks), term (delivery ≥ 37 weeks), and postpartum preeclampsia. This maternal condition is caused by a dysfunctional placenta that secretes factors into the maternal bloodstream, leading to extensive maternal endothelial dysfunction and systemic inflammation^[15]^. Although innate immune cells play a role in supporting the early stages of pregnancy, they also display immunomodulatory properties that are aimed at regulating immune responses against the fetus. Reports indicate that during PE, immune cells accumulate at relatively high densities near spiral uterine arteries and undergo extensive degranulation, which results in the release of substantial amounts of histamines^[16]^.

Compelling evidence implicates maternal innate immunity in PE pathogenesis. Deciphering aberrant signaling networks and effector cell functions may uncover actionable biomarkers and therapeutic vulnerabilities. Macrophages predominantly exhibit the M2 phenotype in a normal physiological pregnancy, whereas there is a shift toward the M1 phenotype in PE^[17]^. M1 macrophages release soluble fms-like tyrosine kinase-1 (sFlt-1), which is linked to impaired angiogenesis in PE^[18]^. Therefore, the shift in the macrophage phenotype from the M2 phenotype to the M1 phenotype indicates a proinflammatory response in PE. The inflammatory response is driven by the release of placental microdebris from syncytiotrophoblasts in PE, with neutrophil extracellular traps (NETs) playing a role in this mechanism^[6]^. Compared with normotensive pregnant women, hypertensive pregnant women present higher circulating NK cell counts, indicating elevated cytolytic activity^[19]^. In rats, the cytolytic activity of NK cells from the placentas of preeclamptic animals was observed to be five times greater than that of sham controls^[20]^.

The complement system, a crucial part of innate immunity, normally shows increased activation during pregnancy^[21]^; however, this activation is significantly amplified in PE^[22]^. Previous studies have demonstrated that activation of the terminal complement pathway, evidenced by elevated levels of the membrane attack complex (C5b-9), is significantly increased in pregnancies complicated by PE compared to normotensive controls^[22]^. T and B cells drive adaptive immune responses that target not only pathogens but also allo- and autoantigens. These responses are marked by the development of immune memory, which strengthens the immune response upon future encounters with the same antigens. It is important to note that PE occurs more frequently in first pregnancies compared to later ones, and the use of barrier contraceptives, which prevent sperm exposure, has been linked to an increased risk of PE^[9]^. To further investigate the connection between PE and immune genes, this study thoroughly examined the expression and regulatory mechanisms of immune-related genes (IRGs) in PE. Additionally, a diagnostic model for IRGs in PE was developed via machine learning techniques.

GO/KEGG enrichment analysis revealed that cytokine−cytokine receptor interactions are involved in PE. Cytokines are glycoproteins or soluble extracellular proteins that serve as key regulators and activators of cells involved in both innate and adaptive immune responses. Various cells in the body typically secrete cytokines in response to specific stimuli, and these cytokines exert their effects by binding to distinct receptors on the surfaces of target cells. Structurally, cytokines can be categorized into different families, and their receptors are similarly classified. Antiendothelial dysfunction peptide in PE (AEDPPE) was found to significantly improve vascular endothelial damage caused by TNF-α and LPS in PE^[23]^. In early-onset PE, vascular endothelial growth Factor A (VEGFA) and VEGF receptor 1 (FLT1) are involved in the cytokine receptor and hypoxia-inducible factor (HIF)-1 pathways^[24]^.

Ensemble machine learning (LASSO coupled with mSVM-RFE) distilled a three-gene signature (SERPINA3, NDRG1, FLT1) from the top eight discriminatory DIRGs. After validation with the test group, we confirmed that the SERPINA3 and NDRG1 genes are associated with PE (AUC>0.7). SERPINA3, also known as Alpha 1-antichymotrypsin (AACT, ACT), is a protein that is part of the protease inhibitor family. It is encoded by the SERPINA3 gene, which is located on the 14q32.13 region of chromosome 14^[25]^. The SERPINA3 protein is also naturally found in the gallbladder, pancreas, testes, and uterus^[26]^. Shortly after placental implantation, there is a transition from early inflammatory Th1 immunity to Th2 anti-inflammatory immune responses^[27]^. The dominant Th2 immunity, which takes precedence over Th1 immunity at the implantation site, safeguards the fetus by balancing Th1 immunity and supporting fetal and placental growth^[27]^. There is an interplay between immunity and inflammation in PE^[28]^. SERPINA3 is classified as a protein involved in acute-phase inflammatory reactions because its serum concentration increases 2–5-fold during immune responses when stimulated by cytokines^[25]^. SERPINA3 is not a primary marker of the inflammatory process; rather, it may reflect the particularly damaging characteristics of that process^[29]^. Significant hypomethylation of the SERPINA3 promoter was detected and identified as a potential binding site for transcription factors involved in developmental processes and stress responses, such as hypoxia and inflammation^[30]^. The expression of SERPINA3N, along with that of IL-6 and TNF, is elevated in all major hypothalamic nuclei due to a high-fat diet and leptin in mice^[31]^. In this study, we further confirmed that the SERPINA3 levels in samples from patients with PE were greater than those in normal samples and that the AUC was greater than 0.79. Our results are consistent with those of previous studies on various diseases related to SERPINA3. IL-6 and its family members can activate the JAK/STAT pathway^[32]^, and SERPINA3 has been recognized as one of the direct transcriptional targets of STAT3^[33]^. These results imply the presence of an IL-6/STAT3 axis in the regulation of SERPINA3 during the inflammatory response; however, the functional importance of this axis in pathological conditions remains unclear.

NDRG1 is a member of the NDRG family, which includes four proteins, NDRG1, NDRG2, NDRG3, and NDRG4, that share 57–65% amino acid sequence identity^[34]^. These proteins are classified within the α / β hydrolase superfamily, even though they lack hydrolytic catalytic activity^[35]^. The NDRG1 gene is located at locus 8q24.2 on chromosome 8. It produces a 2997-bp mRNA, which is then translated into a mature protein with an approximate molecular weight of 43 kDa and consists of 394 amino acids^[36]^. NDRG1 participates in various biological functions and does not appear to play a singular role^[37; 38]^. NDRG1 expression was first detected in the BeWo trophoblast cell line^[39]^. NDRG1 gene mRNA is highly expressed in human placental villous tissue, and its expression is closely related to trophoblast cell damage^[40]^. In human trophoblast cells, NDRG1 is associated with the stress response, proliferation, and differentiation. NDRG1 expression can be induced in primary cultured trophoblast cells under hypoxic and simulated hypoxic conditions^[40]^. NDRG1 expression is increased in placentas from pregnancies affected by severe PE or intrauterine growth restriction (IUGR). This observation indicates a trophoblast response to hypoxic damage^[41]^. NDRG1 inhibits angiogenesis in PE, and the regulation of angiogenesis by NDRG1 may involve the PI3K/AKT signaling pathway^[42]^. Our findings are consistent with those of other studies, indicating that the NDRG1 gene plays a specific role in the pathogenesis of PE.

We applied the CIBERSORT algorithm and reported that Tregs and plasma cells in normal samples were significantly lower than those in PE samples. However, compared with those in PE samples, the numbers of M2 macrophages and monocytes in normal samples were significantly greater. In our study, both NDRG1 and SERPINA3 were negatively correlated with M2 macrophages. Elevated macrophage infiltration coupled with diminished M2 polarization in FGR implicates proinflammatory macrophage subsets in disease pathophysiology^[43]^. These findings suggest that the NDRG1 and SERPINA3 genes may be closely related to the immune system and the pathogenesis of PE.

Several limitations should be acknowledged. Although IHC confirmed SERPINA3 and NDRG1 expression, mechanistic insights would benefit from techniques like flow cytometry. Furthermore, the predictive nomogram requires validation in larger cohorts before clinical translation due to current sample size constraints.

## Conclusion

This multi-omics investigation demonstrates the utility of machine learning approaches in uncovering novel immune biomarkers for preeclampsia, specifically highlighting SERPINA3 and NDRG1. The immune-related genes NDRG1 and SERPINA3 were found to be highly expressed in PE. Moreover, a lower proportion of M2 macrophages and a greater proportion of Tregs were observed in the PE group than in the normotensive group.

## Acknowledgments

”We acknowledge the open-access resources provided by the National Center for Biotechnology Information (GEO) and the National Institute of Allergy and Infectious Diseases (ImmPort).

## Data availability statement

These data were derived from the following resources available in the public domain: The GEO database (https://www.ncbi.nlm.nih.gov/gds), The Import database (https://www.immport.org/shared/),

## Ethics statement

This investigation received ethical approval from the Second Affiliated Hospital of Fujian Medical University Institutional Review Board (No. 2024-321). All procedures adhered to institutional guidelines and national regulations, with written informed consent obtained from every participant.

## Author contributions

Conceptualization, Zhuna Wu and Yumin Ke; Methodology, Weihong Chen and Zhimei Zhou; Software, Yajing Xie; Validation, Zhuna Wu, Shihong Chen and Yumin Ke; Formal Analysis, Zhuna Wu; Investigation, Li Huang; Resources, Liying Sheng; Data Curation, Zhuna Wu; Writing – Original Draft Preparation, Zhuna Wu; Writing – Review & Editing, Zhuna Wu, Shihong Chen and Yumin Ke; Visualization, Yueli Wang and Binbin Chen; Supervision, Congmei Yang and Yumin Ke; Project Administration, Zhuna Wu; Funding Acquisition, Zhuna Wu and Yumin Ke. All authors read and approved the final manuscript.

## Additional files

Table S1. List of Immune-Related Genes from the ImmPort Website.

Supplementary S1: GO analysis results of DIRGs.

Supplementary S2: KEGG analysis results of DIRGs.

